# inSēquio: A Programmable 3D CAD Application for Designing DNA Nanostructures

**DOI:** 10.1101/2024.03.27.586810

**Authors:** Curt LaRock, Paul Sorensen, Douglas Blair, Dabrien Murphy, James O’Connor, Steven Armentrout

## Abstract

DNA nanotechnology is evolving rapidly, paralleling the historic trajectory of the 1970s electronics industry. However, current DNA nanostructure (DN) design software limits users to either manual design with minimal automation or a constrained range of automated designs. inSēquio Design Studio, developed by Parabon^®^ NanoLabs, bridges this gap as a programmable 3D computer-aided design (CAD) application, integrating a domain-specific graphical editor with a Python API for versatile DN design.

Developed in C++ for Windows^®^ and Macintosh^®^ systems, inSēquio features a user-friendly GUI with extensive CAD tools, capable of managing complex designs and offloading computational tasks to the cloud. It supports various DNA design formats, PDB molecule integration, residue modifications, and includes preloaded designs and thorough documentation.

With its combination of features, inSēquio enables a code-centric design (CCD) approach, enhancing DN construction with improved precision, scalability, and efficiency. This approach is elucidated through a streptavidin barrel cage designed via Python notebook and a spheroid origami case study.

Marking a significant advance in DN design automation, inSēquio, the first fully programmable 3D CAD tool for DN design, enables both manual and programmatic 3D editing. This fusion of features establishes inSēquio as a transformative tool, poised to significantly enhance designer productivity and expand the scope of possible designs.

**Extended Abstract:** Advances in DNA nanotechnology have positioned the field at a juncture reminiscent of the pivotal growth phase of the electronics industry in the 1970s. The evolution of software for designing DNA nanostructures (DNs) is following a similar historical trajectory and dozens of software packages have been developed for creating them. Existing software options, however, require users to choose between manual design with minimal automation support or selecting from a limited set of designs, typically wireframe, that can be generated from a high-level structural description. Here, we introduce the inSēquio Design Studio, a programmable 3D computer-aided design (CAD) application that effectively bridges this gap. By integrating a domain-specific, freeform graphical editor with a Python application programming interface (API), inSēquio provides a comprehensive and extensible platform for designing complex nucleic acid (NA) nanostructures.

The inSēquio desktop application, developed in C++, runs on Windows^®^ and Macintosh^®^ operating systems. Its graphical user interface (GUI) features multiple synchronized view panels and a diverse set of CAD and NA-specific editing tools. Its optimized graphics pipeline enables editing of designs with >2M nucleotides, and it includes an integrated service infrastructure for offloading heavy computations to cloud servers. The software also supports import and export of various DNA design file formats, integration of arbitrary PDB molecules, and specification of residue modifications. Additionally, it includes preloaded sample designs, scripts, and comprehensive documentation.

Parabon has used evolving versions of inSēquio for over a decade to design a variety of proprietary DNs and have now transitioned it into a commercially available product. This paper summarizes inSēquio’s features, discusses its strengths and limitations, and outlines planned enhancements. Although freeform 3D design is well supported in inSēquio, the integration of its CAD environment with its API facilitates a *code-centric design* (CCD) approach for DN construction that offers notable productivity advantages over traditional methods, including enhanced precision, scalability, and efficiency. Here we describe CCD, outline its benefits and demonstrate its use through a well-documented Python notebook, included with the product, which generates a sample design within the inSēquio application. A spheroid origami created using CCD is also presented.

As the first commercial fully programmable 3D CAD application specifically created for DN design, the release of inSēquio represents a milestone in the field of DN design automation. It introduces a new dimension to the discipline by enabling both manual and programmatic 3D editing, thereby facilitating an innovative CCD approach. The availability of extensive documentation and technical support enables designers to efficiently adopt and utilize these capabilities. This combination of features establishes inSēquio as a noteworthy addition to the tools available for DN design, with the potential to significantly increase designer productivity and broaden the scope of designs that can be developed by practitioners of all skill levels.

Windows and Mac versions of the inSēquio desktop application are available for download at https://parabon.com/insequio.

**Graphical Abstract:** 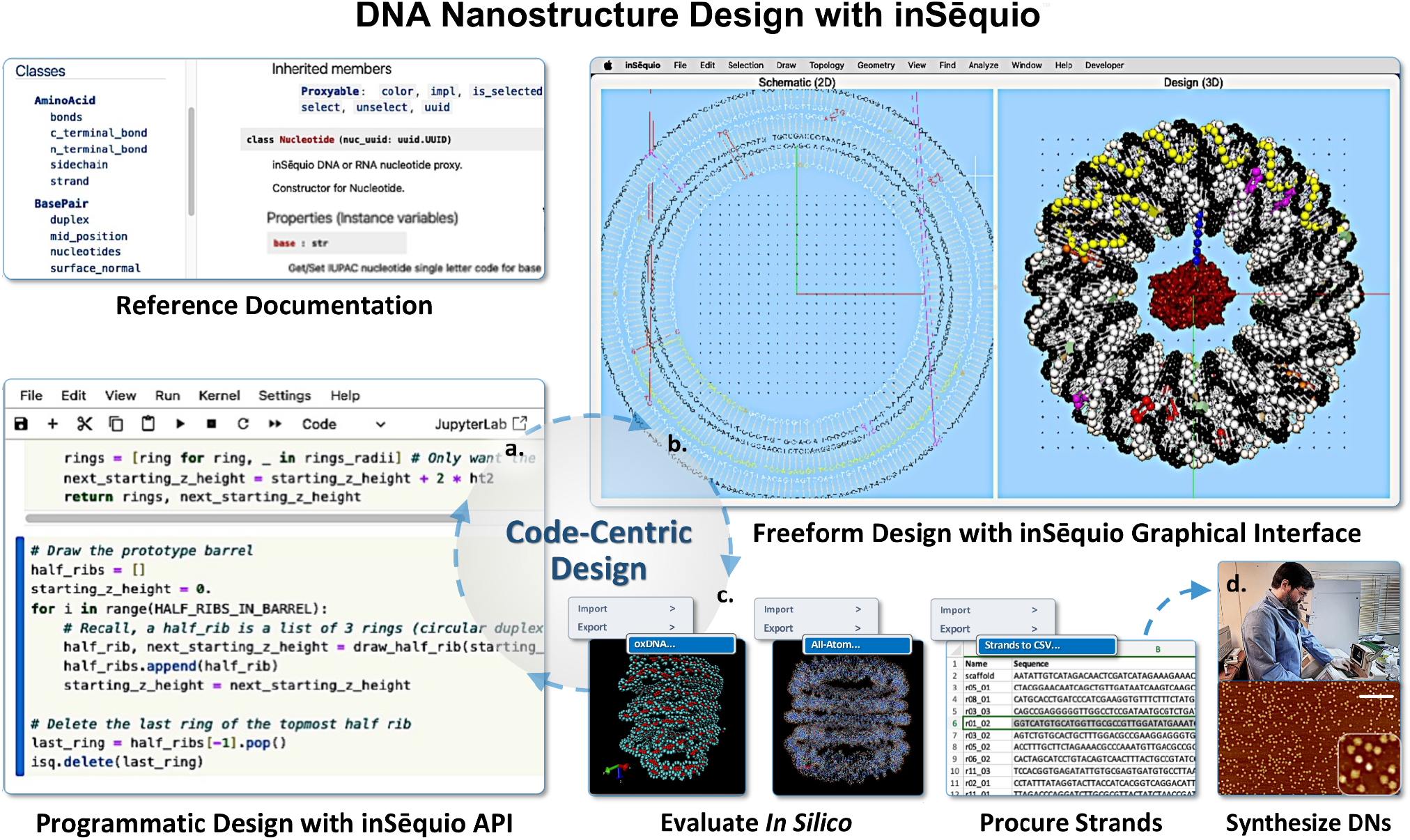

An illustration of the inSēquio Design Studio desktop application interoperating with a Python Jupyter notebook and molecular dynamics (MD) simulation tools to support an iterative code-centric design (CCD) process. The design cycle includes **(a)** programmatic and/or manual creation of objects in the inSēquio editors; **(b)** visual inspection and manipulation of objects via user interface; **(c)** *in silico* evaluation of designs via MD simulation using native or external tools; repeating **a-c** as necessary; and **(d)** procurement of strands and synthesis of DNA nanostructures (DNs).

## INTRODUCTION

In fields as diverse as naval architecture and microelectronics, the challenges of product design have spurred development of computer-aided design (CAD) software to manage complexity and streamline design workflows. The significance of design software for the field of DNA nanotechnology was evident from the early work of its founding father, Nadrian Seeman^1^. Our own interest in such tools stemmed from his personal demonstration of GIDEON, arguably the first 3D DNA CAD program, just when it was becoming operational^2^. Paul Rothemund’s introduction of DNA origami cemented software’s critical role in the field^3^.

The evolution of DNA design packages since that time has been well covered in recent reviews^4,5,6^. Dey et al. classified DNA design software into three generations, roughly characterized as G1) graphical editors (e.g., cadnano^7^ and Tiamat^8^), G2) top-down wireframe generators (e.g., vHelix^9^, DAEDALUS^10^, etc.) and G3) editors that also integrate second-generation features and simulation tools (e.g., Adenita^11^, ATHENA^12^).

oxView, ENSnano and MagicDNA represent the latest third-generation tools. oxView is a browser-based visualizer that provides robust simulation and analytical tools; however, the authors indicate it is not intended for de novo design, despite possessing basic editing functions^13,14^. ENSnano provides 2D editing in a split view to facilitate 3D design, and can display resulting 3D designs, but has no 3D editing features besides the ability to make a crossover^15^. MagicDNA is a collection of MATLAB-based CAD tools for producing conventionally shaped components – wireframe, surface, and lattice structures – that can be connected to form larger DNA assemblies^16^. The latest release moves closer towards genuine freeform design by adding tools for converting splines into curvilinear DNA bundles^17^.

Catana offers tools for combining DNs with proteins; however, it has few DNA editing capabilities and expects designs to be imported^18^. The Common-Lisp framework ‘small’ emphasizes the importance of programmatic design, but lacks a graphical editor and depends on external tools for visualization^19^. DNAxiS is a collection of Python algorithms for generating designs with axial or cyclically symmetric curvature that can be displayed via its web server^20^.

These and other software contributions have facilitated creation of a diverse array of DN designs. It is evident, however, though not unexpected, that advancing CAD editing interfaces has not been a research priority. Nonetheless, the evolution of design needs compels us to look beyond conventional improvements. It is well recognized that even the most sophisticated CAD user interfaces (UIs) ultimately encounter their limits in effectively managing complexity and ensuring precise control — requirements that invariably demand *programmability*. We suggest that CAD editors specifically tailored for nucleic acid (NA) design, when complemented by an associated API, represent what might be termed fourth-generation (4G) DN design environments.

An example 4G environment from the literature is scadnano. Drawing inspiration from the graphical UI design of cadnano – the most popular CAD application in the field – scadnano (“scriptable cadnano”) offers a comprehensive Python API to augment its graphical editing capabilities^21^. However, like cadnano, scadnano is limited to 2D editing of designs, requiring one to export to the oxDNA^22^ format to visualize the 3D structure in oxView. Despite this limitation, it exemplifies the advantages of combining manual with programmatic editing for DN design.

4G design environments, which combine both manual and programmatic control over the design process, offer a blend of advantages not achievable by either approach alone. Programmatic control facilitates automation of repetitive tasks, algorithmic optimization of design topology and geometry, and enhanced reproducibility and adaptability, permitting swift modifications to design features without the tedious overhead of editor-only design. Conversely, manual UIs provide the ability to rapidly prototype design concepts, make modifications that are difficult or costly to codify, and an interactive environment for creative exploration. Together, these dual capabilities facilitate a *code-centric design* (CCD) approach to the challenging task of DN design that provides practitioners significant gains in efficiency and design quality. In this paper, we introduce our 4G contribution, the inSēquio Design Studio, the first fully programmable 3D CAD editor created exclusively for designing DNA and RNA nanostructures. (For brevity, our discussion hereafter refers only to DNA nanostructures [DNs].) This paper provides an overview of inSēquio’s primary and differentiating features, but does not comprehensively describe the product. Detailed documentation, tutorials, and the Python API are available for review with the free trial version of the software available at https://parabon.com/insequio.

## MATERIALS AND METHODS

### Software development objectives and system requirements

Whereas the initial implementation of inSēquio (not released) was limited to 2D schematic editing, the current version (v1.0.4.3) was architected as a 3D CAD application from its inception. Recognizing that nanoengineers might have limited experience with CAD applications, we aimed to blend the 3D object manipulation capabilities essential for CAD design with drawing tools and functions typically encountered in standard office software suites. We opted for desktop versus browser-based implementation to maximize UI richness and responsiveness, and to grant users direct control over data security - designs stay local unless deliberately transmitted for server-based processing. This also allowed us to implement our own graphics pipeline and optimize its performance to enable extended reality product extensions.

The Qt application framework was chosen to facilitate cross-platform development. This release of inSēquio is compatible with Windows^®^ 10 and 11 (x86-64 only) and Macintosh^®^ macOS 12/Monterey and later (Intel and Apple Silicon). We anticipate releasing a version for Linux in the near future. We chose Python (v3.10 or later) for the initial API implementation because of its pervasive use for interactive scientific computing. While extremely large designs benefit from computers with large RAM, we routinely run inSēquio on laptops with 16GB of RAM and we expect good performance on less capable computers.

## RESULTS

### The inSēquio desktop application

#### Synchronized view panels

inSēquio’s main application window has three view panels: Schematic (2D), Design (3D) and Analysis (3D). Users can create and edit objects in the Schematic and Design views, either manually or through the API. When an object is created using the API, its representation can be directed to appear in either view or both. In cases where the object is generated in both views, their representations will be synchronized, i.e., selections and many types of edits made in one view will automatically be reflected in the other (**Figure 1**). Objects resulting from various analyses are generated in the Analysis view and are synchronized with their corresponding elements in the other views. This integration of different yet synchronized views has, in our experience, offered unique design insights and significantly enhanced the detection of potential issues that could otherwise be easily overlooked.

**Figure 1.**
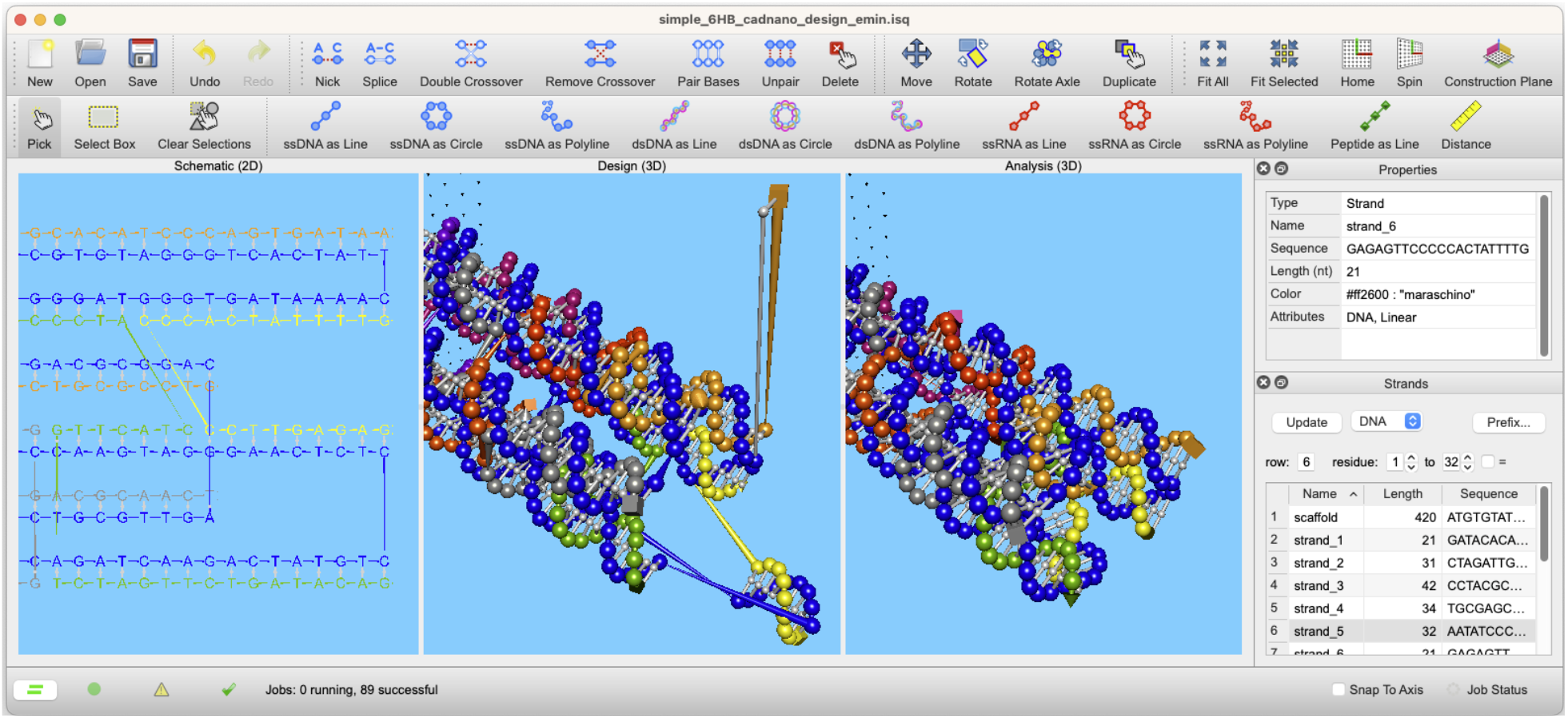
A simple six-helix bundle imported from cadnano, a portion of which is shown in all three view panels. A nucleotide and duplex have been deliberately repositioned in the Design view leaving them with unnaturally long bonds. The Analysis view shows the result of performing energy minimization on contents of the Design view calculated using the **Minimize Energy** command, which eliminates the long bonds. Selection of an object in any one view highlights it (in yellow by default) in all synchronized views as shown.

#### Native molecular species

DNA and RNA nucleotides and amino acid (AA) residues are represented as visually distinct single-particle beads that can be drawn individually or as linear, circular or polyline strands and duplexes (nucleic acids only). NA strand termini are distinguishable by their geometric forms, specifically, 5’ and 3’ nucleotides are represented as a cube and cone, respectively. Likewise, peptide N- and C-termini are represented by rectangular and triangular pyramids, respectively. A rich set of G-quadruplexes is also provided.

To facilitate functionalization, residues can be decorated with chemical modifications selected from a built-in catalog; users can also create their own modifications for inclusion in the catalog. Modification codes are persisted with strands when they are exported to CSV file format (typically for procurement). A generic linker object can also be used to connect species in non-standard manners, such as splicing the 5’ ends of two NA or joining a peptide to a NA. (planned enhancements for this feature are discussed in Future Directions).

Object mesh representations of other molecule types can also be drawn. A small set of molecules is included and users can install others (e.g., from PDB) using the **File** → **Import** → **Molecule Mesh** menu. **Figure 2** shows how these species appear in inSēquio’s Design view.

**Figure 2.**
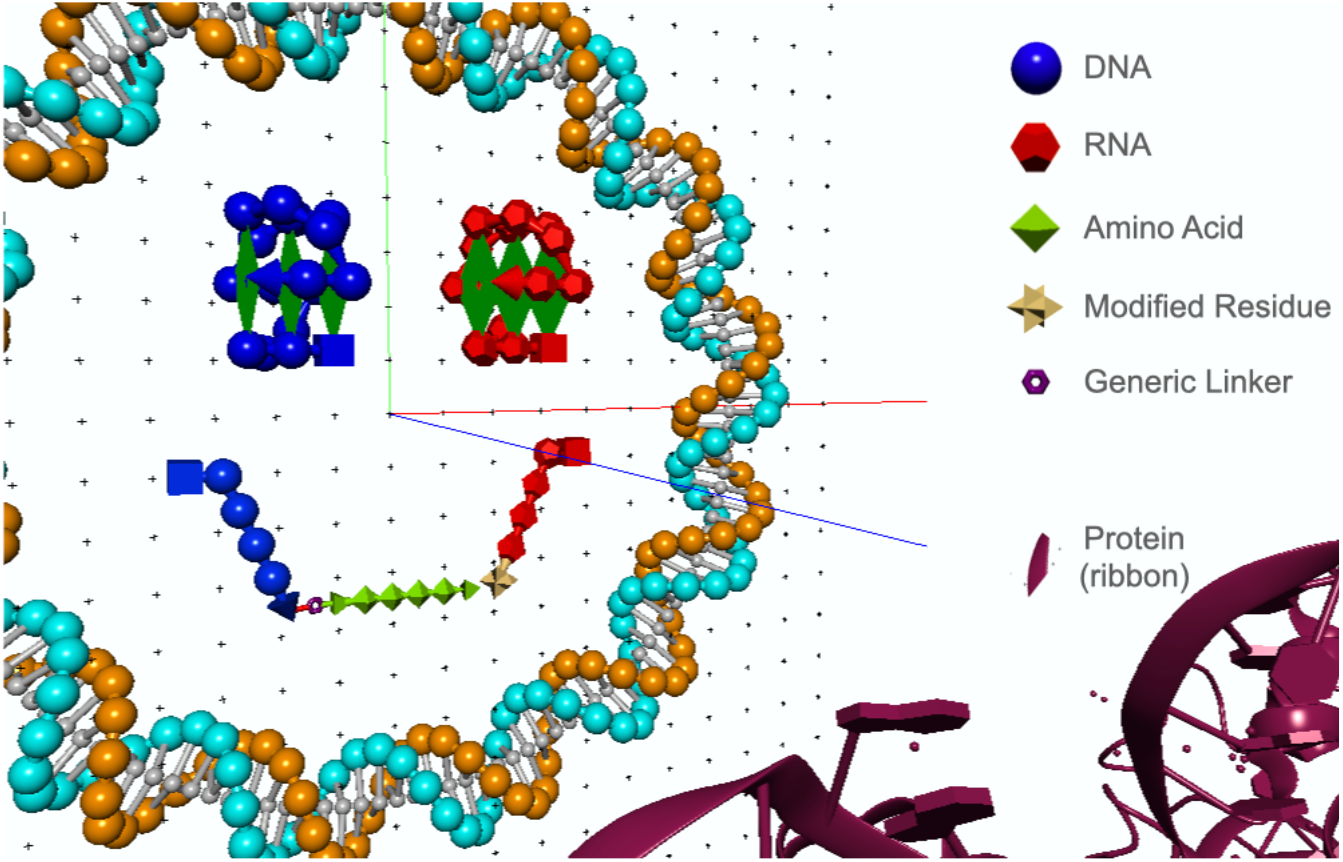
The molecular species natively supported by inSēquio. Circular DNA duplex (cyan and orange) surrounds a DNA parallel G-quadruplex (blue) and RNA antiparallel G-quadruplex (red), each comprised of three tetrads (four nucleotides connected by green rectangles). Single-stranded DNA (blue) is conjugated to a short peptide (lime) via a generic linker object with user defined connections. A single-stranded RNA strand (red) has a 3’ terminal that is modified with 3’ biotin-TEG (gold stellated octahedron). A portion of a protein, imported as a ribbon mesh, is in the foreground (maroon).

### Freeform editing in 3D

Navigating three dimensions in CAD applications presents unique challenges, particularly given the 2D constraints of standard mouse and keyboard interfaces. inSēquio addresses these challenges with a rich set of inherently 3D CAD editing features. Central to the 3D interface is the application’s Construction Plane (CP), which can be positioned anywhere in the 3D Design view via intuitive six-degrees-of-freedom (6DF) controls. The CP panel also offers customization options for its appearance, such as adjusting its axis visibility, grid extent, and grid density. Object creation, movement and rotation are performed with respect to the current orientation of the CP. Objects can also be moved by arbitrary distances and angles via the **Geometry** → **Transform Selected** command, the UI panel for which provides convenient 6DF controls.

Commands for topologically editing strands, found in the **Topology** menu, include **Nick** (cut a strand), **Splice** (join two strands), and **Pair** and **Unpair** nucleotide bases. The **Cross** command performs the nicking and splicing commands to effect a double-crossover; **Uncross** inverts these operations. Geometric mouse editing commands include **Move**, **Rotate**, and **Rotate Axle**; these actions can also be effected precisely with the **Transform Selected** command, which accepts six conventional spatial transformation parameters. A variety of other editing commands are provided for manipulating strands and their sequences (**Figure 3**). The design in **Figure 4** demonstrates what one designer accomplished using these freeform editing tools in ∼10 minutes at the end of a long day.

**Figure 3.**
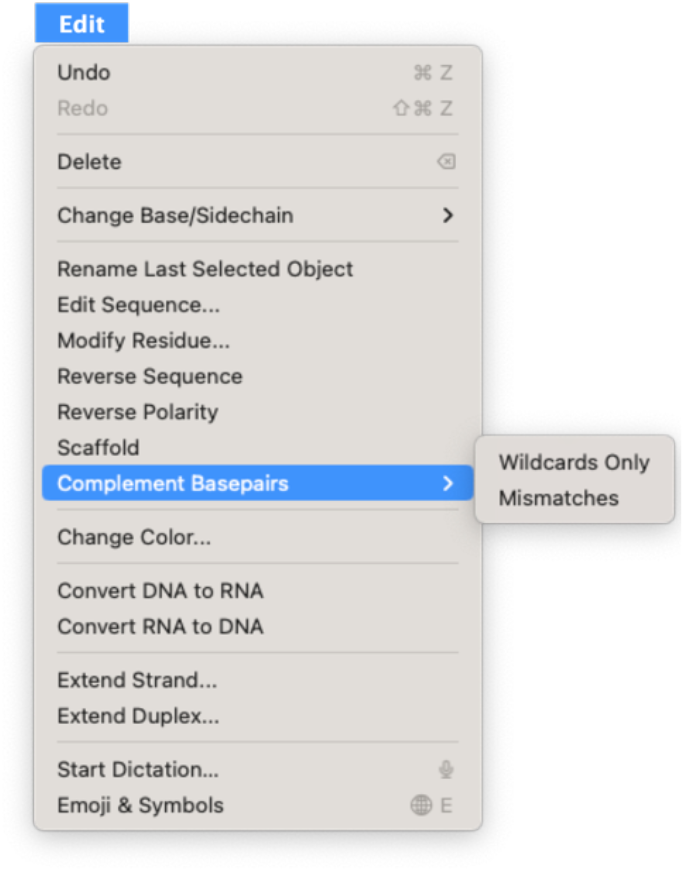
The Edit menu provides several commands for manipulating strands and sequences.

**Figure 4.**
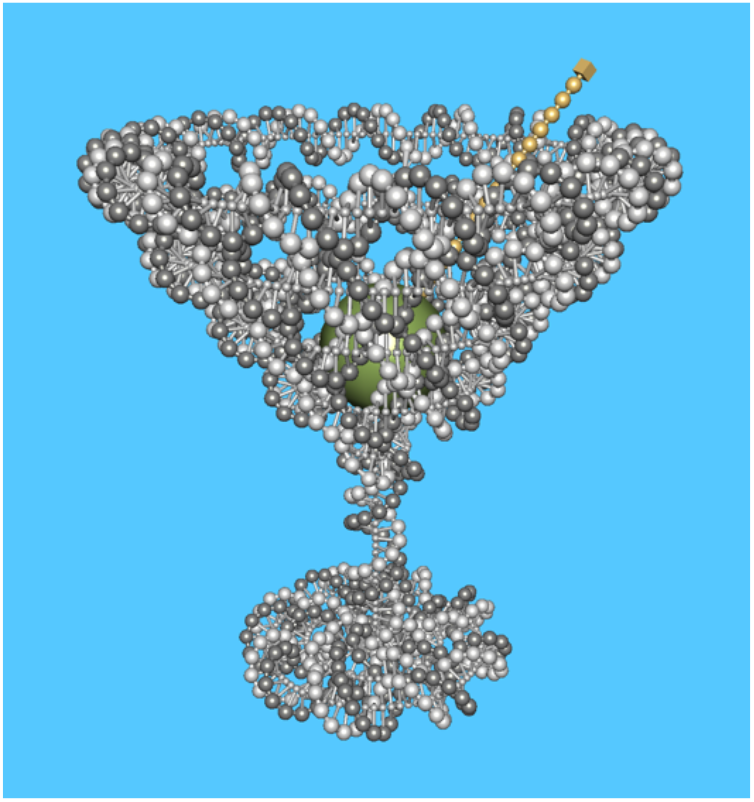
A playful design created entirely with 3D freeform editing tools.

### Data import and export

The current version of inSēquio is a single-document application, hence Copy/Paste between two inSēquio documents is not possible. However, structures from multiple inSēquio files can be combined by importing a source design into a target design using the **Import** command. cadnano files can also be imported, creating both 2D and 3D representations in the Schematic and Design views, respectively. NA components of a design can be exported to oxDNA^22^ and PDB^23^ file formats. inSēquio uses oxDNA^24^ and tacoxDNA^25^ software components in the implementation of its import and export services.

Standard-resolution images, as well as high-resolution images suitable for publication, can be copied to a file or the clipboard using the **Capture Image** command. The former is a pixel-for-pixel reproduction of the designated view; the latter produces the same image at 5x resolution.

#### Functionalization via residue modifications and generic linkers

The application potential of DNs arises from their capacity for molecular species functionalization. They uniquely enable colocation, spatial organization, and transport facilitation for a wide range of molecules, distinguishing them from other current nanoengineering technologies. A variety of oligonucleotide modifications are now available to facilitate conjugation with peptides, antibodies, small molecules and many other species. To support inclusion of such modifications in NA designs, inSēquio has a **Modify Residue** command that displays a catalog of standard oligo modifications (e.g., 5’ Hexynyl, 3’ Cholesterol-TEG and Cy5 fluorophore among many others). Entries include customizable fields such as the text code used during export of strand sequences to CSV format (e.g., `/5Hexynyl/ACGTACGT`), an image of the chemical structure, detailed notes (e.g., from the vendor), SMILES codes and references. Users can simultaneously apply modifications to one or more selected residues after which their shape changes to a stellated octahedron to facilitate easy identification. Additionally, the **Draw** → **Linker** command can be used to insert a namable and colorable hexagonal ring that can be spliced to strand termini to represent a generic conjugation between strands.

#### Find commands and metrics

The **Find** menu contains 24 entries for locating various components of a design. It includes a comprehensive **Sequence…** dialog for specifying the type and composition of search probes, entries for locating problematic regions of a design (e.g., strands that are too long or duplexes that are too short), nucleotides with various properties (e.g., paired, unpaired, having wildcard bases, bulge nucleotides, etc.), strands with various properties (e.g., circular strands, strands marked as scaffolds) and other structural elements, such as duplexes and nicks. A list of design level properties can be generated using the **Design Metrics** command and distances between objects in any view can be displayed with the **Measure** tool.

#### Software architecture

The inSēquio architecture consists of a set of software components specifically designed to efficiently capture, represent, and render DN designs, as detailed in **Figure 5**. The **Object Model** captures the essential elements of a design, including nucleotides, strands, duplexes, amino acids, peptides, modifications, linkers, and other molecules, along with their corresponding topology. The **Representation** module maps topology to 2D and 3D geometry. The **Scene** module manages the presentation of this representation to the user as a 2D or 3D scene, incorporating aspects like material, lighting, and camera position. The **Renderer** then converts these 3D scenes to 2D images for display on the user’s screen. The **Internal inSēquio API** exposes operations for manipulating the Object Model, its Representation, and the Scene. The **Graphical User Interface** (GUI) allows users to create, modify, view, and analyze designs, with UI operations implemented via calls to the Internal inSēquio API and results rendered by the Renderer.

**Figure 5.**
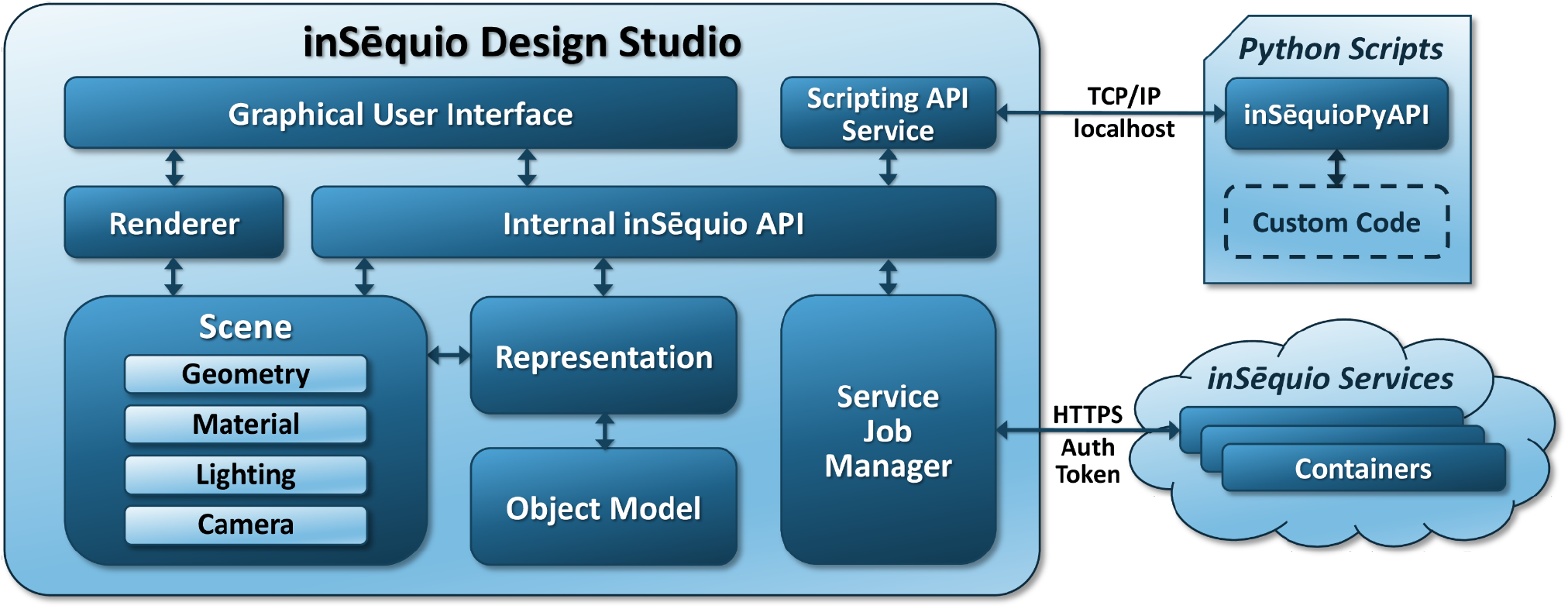
Architectural diagram of inSēquio illustrating its key components and their interconnections.

A distinguishing characteristic of the inSēquio architecture is its support for both manual and programmatic design: users can engage with the GUI for interactive editing and/or utilize the intuitive and flexible inSēquio Python API (**inSēquioPyAPI**) for automated, script-driven design. The implementation of the underlying Python API connects to the inSēquio application through a localhost TCP/IP connection, exchanging JSON-encoded commands and responses. The **Scripting API Service**, which listens to connections on a configurable port number, processes these commands via the Internal inSēquio API and returns corresponding responses.

To maximize scalability and performance, the inSēquio Design Studio application was implemented in C++ and OpenGL, which provides a standardized way to implement GPU-accelerated graphics. Its graphics pipeline employs highly-optimized custom shaders and scene caching to achieve a native refresh rate greater than 90Hz, which is essential for enabling freeform design, editing, and visualization in extended reality (XR; virtual and augmented) environments. These capabilities were implemented in inSēquio for Oculus Rift, as demonstrated at MADNano 2019^26^ and FNANO 2021^27^, and we anticipate product extensions to enable XR use of inSēquio in the future.

On contemporary laptop computers, this design enables construction of DN designs comprising over two million nucleotides (**Figure 6**). Additionally, to support computationally intensive operations like energy minimization and calculating oxDNA trajectories, we developed a backend service infrastructure and a REST-based protocol for remote execution of computational jobs initiated by inSēquio commands. This service architecture allows users to perform analyses that may be too demanding for local hardware. User requests are managed by the **Service Job Manager** (SJM), which authenticates to **inSēquio Services** using the user’s credentials to obtain an authentication token for subsequent interactions. All communications between the application and inSēquio Services are secured using Hypertext Transfer Protocol Secure (HTTPS). The SJM sends the user’s request to inSēquio Services, which then runs the request asynchronously in a designated Docker container. The SJM monitors the job and notifies the user upon completion. Users can then view the results through the SJM interface.

**Figure 6.**
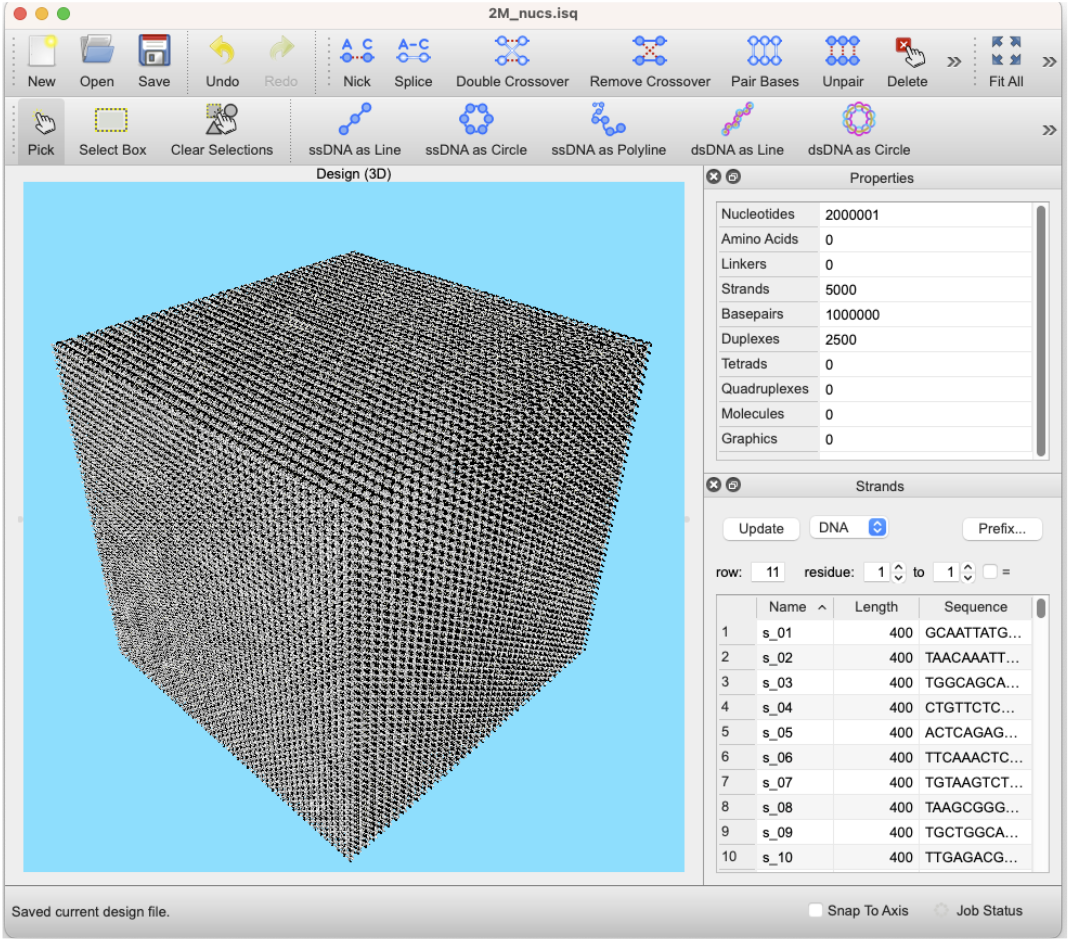
A cube containing >2 million nucleotides (2500 staples).

#### Data Security and Privacy

The inSēquio Design Studio and inSēquio Services were designed to maintain the security and privacy of user data. User data stays on the local machine, except when executing authorized inSēquio Service operations, such as performing an energy minimization. Before transmission to inSēquio Servers, these operations alert users of pending data transfers, offering an option to cancel. During transit, data security and integrity are safeguarded using HTTPS.

On inSēquio Servers, several measures ensure data protection. Data at rest are secured on encrypted drives. Processing occurs in isolated Docker containers to guarantee customer data are not commingled. Access to inSēquio Services and Servers requires authentication, with robust access controls in place to prevent unauthorized access. Parabon staff access to inSēquio Servers is rare and only occurs under specific circumstances, such as to address technical issues. Such access is governed by strict protocols, ensuring it is minimal and directly related to problem resolution. Robust audit mechanisms are employed to enforce these protocols and guarantee the integrity and confidentiality of user data. Finally, inSēquio Servers persist user data only for the duration of an active job; upon a user’s deletion of the job, all associated data are immediately and completely erased.

#### Documentation and Technical Support

A key differentiator for inSēquio is its robust **Help** system, which includes a comprehensive set of user support resources to assist designers in both learning and effectively utilizing the software. The **User Guide** details every aspect of the interface, supplemented with Tutorial videos and a “Cookbook” that elucidates complex operations. Keyboard shortcuts and mouse and trackpad actions are also detailed within a **Keyboard Shortcuts** window. The Python API, described in the next section, is equally well-documented.

A significant advantage of inSēquio’s commercial support is its team of dedicated developers and staff who offer prompt and thorough technical assistance. This ensures designers receive the help they need to fully utilize the software and accelerate their research. For those requiring design assistance beyond standard technical support, consulting services are available.

### The inSēquio Python API

While the inSēquio GUI offers an intuitive and robust platform for freeform DN design, its Python API significantly enhances the software’s capabilities. It allows for direct programmatic construction and editing of DNs within the application’s 2D and 3D graphical editors, making these structures immediately editable and interactively manageable, a novel capability in the field. By enabling designers to blend script execution with GUI-based inspection and manipulation, this integration of API and 3D CAD interface not only enhances control and visibility within the design process but also overcomes the limitations of traditional manual design tools and automated ’black box’ design generators, marking a significant advance in DN design capability. Having established the synergistic benefits of integrating API and CAD for DN design, we now describe the specific molecular classes within inSēquio that facilitate this advanced design approach.

#### Molecular classes

The primary Python classes in the inSēquio API represent molecular species and their subcomponents. These classes, defined in the insequio.structure module, all inherit from the abstract class `Proxyable`, which functions as a hidden proxy, discreetly facilitating communication with the application’s **Object Model**. The class hierarchy deriving from `Proxyable` (**Supplemental Figure S1**) includes abstractions for `Strand`, made up of `Residues` connected by `Bonds` that, along with a `Nucleotide` subclass of `Residue` for nucleic acids, and the `AminoAcid` subclass of `Residue` for amino acids, collectively support concrete representations of DNA, RNA strands, and polypeptide chains. `Nucleotide` objects can be paired to form `Basepair` objects that, when arrayed, comprise `Duplex` objects.

Nucleotide bases and amino acid side chains can be variably specified using IUPAC characters; for instance, nucleotides can have a base value of “N” (wildcard) or “R” (“puRine” – A or G) in addition to “A”, “C”, “G” or “T.”

All derivatives of `Proxyable` can be named and colored. `Residue` and `Basepair` objects are positioned in 3D, and optionally 2D, using `Point3` and `Point2` objects, respectively, which are facades for NumPy points. While methods like `dna_duplex_line` generate objects with plausible geometry, the system allows creation of implausible structures (e.g., overly distant nucleotides connected by an unrealistically long bond). Rigid body constraints are not enforced, permitting object intersections during construction. We have found this flexibility to be generally useful, dampening the motivation to implement code to enforce physical constraints, especially since the **Energy Minimization** command can be used post construction to eliminate such occurrences.

#### API and object model interaction

API sessions begin with a call that establishes a TCP connection between the Python interpreter and the inSēquio application. Once this connection is established, users can make API calls to edit or access the application’s current design. Importantly, the API does not strive to maintain a separate, synchronized version of a design in memory. Instead, Python objects that represent design components merely act as references to the C++ object model. The code in **Figure 7** illustrates this point. It creates a linear DNA duplex 10 nm in length and renders it in the inSēquio application before printing information to the developer’s console. When the `num_basepairs` property of `duplex` is accessed, it retrieves the current value from inSēquio’s object model each time it is called, rather than merely accessing an attribute of the Python variable `duplex`. This deliberate *separation of concerns* between the API and design representation reduces system complexity thereby improving the maintainability of both application and API code bases. It also enables designers to seamlessly combine both manual and programmatic editing within a single session. In fact, users can interact with the inSēquio application, for instance rotating objects in the Design view, even while API code is actively editing a design (e.g., during a long-running block of code).

**Figure 7.**
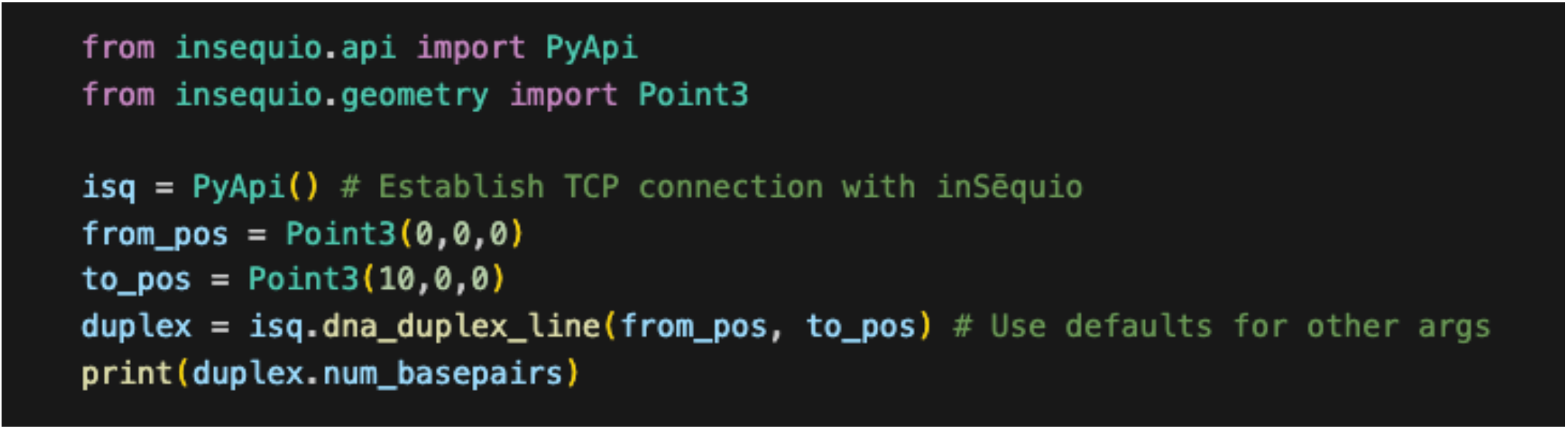
A code snippet that creates a linear DNA duplex in the inSēquio Design view and then prints the number of base pairs it contains to the developer console. The value for the `num_basepairs` property is not stored with the Python object `duplex`, but is fetched from inSēquio’s C++ object model.

#### Integration with the inSēquio UI

Python scripts, developed using the inSēquio API, can be launched from the inSēquio Developer menu. This feature streamlines the creation of custom automation tools for a wide array of design and analysis tasks. Such integration embeds the scripts into the inSēquio UI, effectively making them built-in features.

These scripts are valuable due to their dual capacity for dialogue-based user interaction and real-time interaction with the displayed design. The inSēquio API facilitates the inclusion of UI controls, such as text and spin boxes, check and radio buttons, dropdown menus, file pickers, and layout widgets within scripts. This empowers script developers to design dynamic user interfaces that manage the collection and processing of analysis parameters, execution of complex tasks, and visualization of outcomes.

Additionally, these scripts can interact with and modify the current design using inSēquio’s robust design interrogation and manipulation features, allowing them to query the state of the current design (e.g., identifying selected design elements) and perform desired analyses and transformations. Scripts may be compiled into bespoke libraries tailored to individual users or domain requirements, optimizing inSēquio’s utility for particular workflows and significantly enhancing both productivity and design quality.

### Code-centric design with inSēquio

#### Foundational concepts

The *code-centric design* (CCD) approach combines code-driven construction, analysis, and optimization with interactive 3D visualization and evaluation, forming an iterative and cohesive method for the efficient development of DNs. Design cycles typically follow four key stages: establishing goals, prototyping the next development phase using GUI or code, conducting visual inspections, and analyzing the results, with the insights gained informing the next cycle. With inSēquio, CCD offers a precise yet flexible approach for constructing DNs that overcomes the limitations of manual design, while still enabling direct GUI-based visualization, editing and manipulation by designers. The advantages of CCD are multifaceted: it allows designers to automate repetitive tasks and manage intricate design elements with high precision and efficiency. This includes scalability for large-scale designs, parameterizability for the rapid creation of design variants, and optimizability for algorithmically refining design choices. CCD represents a paradigm shift in DN design, blending the rigor of computational logic with the creativity of freeform design. In this way, CCD shares similarities with the design process in computer-animated film creation, where programming and software tools play a central role in bringing animated narratives to life.

Generally, CCD is an iterative process that involves incrementally developing and running code snippets for each design step, followed by observing and evaluating the outcomes in the graphical editors. Computational notebooks, with their interactive execution environment, provide an ideal complementary platform for employing CCD with inSēquio, enabling users to iteratively develop, visualize, and fine-tune designs in real-time. Each iteration typically begins anew, progressing from a blank document to the most recently completed design stage. This cyclical approach, coupled with the interactive investigation capabilities of the inSēquio application, enables continuous improvement and fine-tuning at each design step, which is essential for producing high quality DNs.

At any point, DN designs can be evaluated through inSēquio’s analysis tools or by observing energy minimization results. Additionally, coarse-grained simulations of designs using oxDNA^24^ can be automatically generated using inSēquio Services via the **Compute oxDNA Trajectory…** command. For in-depth analysis, designs can be exported to all-atom formats for molecular dynamics simulation (although not yet as an inSēquio Service). These simulations are valuable as they often reveal opportunities for design refinement, where the true efficiency of CCD becomes evident. For critical design elements, such as an origami scaffold route, modifications are more efficiently executed when these elements are initially created through code. This efficiency proves particularly valuable in scenarios often encountered in practice, where extensive modifications or a complete reevaluation of strategy are necessitated by the discovery of design flaws. In such cases, straightforward code adjustments or parameter tweaks can significantly improve the design, thereby saving substantial time compared to the effort required for manual redesign.

#### Algorithmic design evaluation and decision-making

One of the most valuable features of inSēquio’s API is its facilitation of computer-aided engineering (CAE) methods, enabling the programmatic manipulation of 3D designs and the analysis of resultant effects. This capability fosters the development of sophisticated algorithms that can explore design space and evaluate alternative design decisions based on their varied consequences, geometric, topological, or otherwise. Examples of these algorithmic applications include identifying candidate crossover locations, selecting optimal crossover networks from these candidates, determining the best relative rotations of neighboring helices, and choosing nick locations for functionalizable overhangs. Notably, tasks like selecting among candidate crossover and nick locations are computationally intensive, with both problems being NP-complete. While we provide only simple search algorithms for these challenges in the notebook presented below, the API has allowed us to create sophisticated algorithms for crossover and nick selection that will soon be incorporated into both the inSēquio API and the application, which will further enhance the tool’s utility for DN design.

#### Code-centric design of a barrel-shaped streptavidin cage

Here we present highlights from a Jupyter notebook, included with inSēquio, that provides users with a richly documented, step-by-step illustration of how the inSēquio API can be used to create a design from scratch, specifically a barrel-shaped streptavidin cage. To facilitate use of the API, the inSēquio application includes a user-friendly setup wizard, **Developer** → **QuickStart**, that creates a Python virtual environment, installs required Python modules and walks users through a ‘Hello World’ script to ensure the Python environment is installed correctly. It then presents the **sample_SA_barrel.ipynb** notebook. Automation of these steps via **QuickStart** eliminates potentially troubling installation issues and thus allows even users with no Python experience to quickly exercise the API.

Using the CCD approach, the major steps for creating a DN, in their recommended order of execution, are: drawing duplexes that form the desired DN shape with topological neighbors close enough for crossovers (helical centers ∼2.7 nm apart); identifying nucleotide quartets for candidate crossovers; selecting a set of candidates that achieve desired crossover density with a minimum spacing between them of ≥ 7 base pairs; making the crossovers at these selected locations; identifying candidate staple nicking sites preferably ≥ 8 nucleotides from any crossover; choosing a set of nick locations that result in staples of desired lengths, typically 17– 60 nucleotides; and finally nicking the staples.

The sample notebook provides code for executing each of these design steps. Users are instructed to successively execute notebook cells and observe the results in the inSēquio graphics editors. As in normal design practice, color is used extensively to illustrate the results of various calculations. **Figure 8** shows the barrel design at three stages of completion.

**Figure 8.**
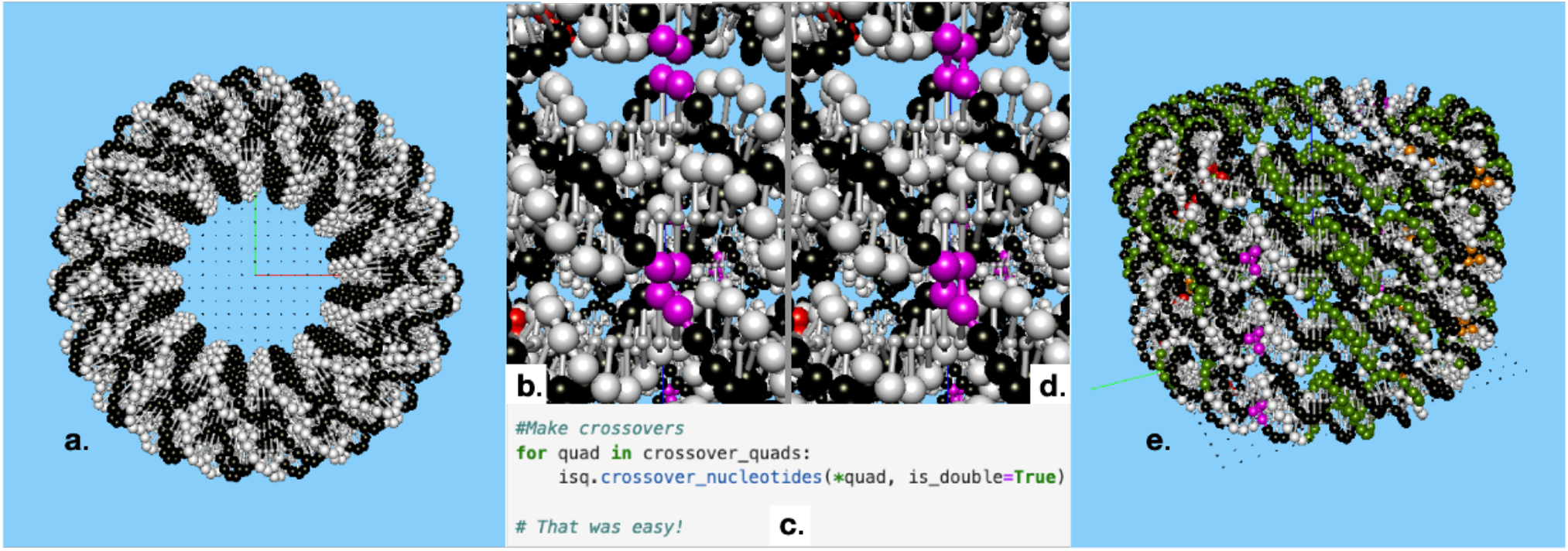
Sample barrel at various stages of design. (**a**) The initial prototype comprised entirely of disconnected circular duplexes each drawn with API method `dna_duplex_circle`; (**b**) Closeup of two quartets of nucleotides (magenta) that have been algorithmically identified as suitable sites for a double-crossover junction; (**c**) The API call to create crossovers with such quartets (`quads`); (**d**) Resulting double-crossovers; (**e**) Candidate nick location sites (green) that are sufficiently distant from any crossover.

The notebook highlights a crucial aspect of API-driven design: edits that modify strand definitions, such as nicking and splicing, invalidate previous strand references (e.g., if two strands are spliced, neither of the original strand references exist afterward). Therefore, it is highly recommended to complete all preliminary decisions for a given design step, such as determining crossover locations, prior to executing these edits through the API. This approach avoids the need to re-identify strand references before proceeding with further calculations or edits.

The barrel notebook showcases a distinctive feature of inSēquio: the capability to concurrently create 2D and 3D versions of a design that are synchronized (for instance, selecting an element in one view automatically selects it in all synchronized views). This synchronization is achieved by providing both 2D and 3D position arguments to drawing functions like `dna_duplex_circle`. This dual-view approach proves to be particularly effective in identifying design defects, as the 2D and 3D perspectives each reveal different aspects of a design (**Figure 9**).

**Figure 9.**
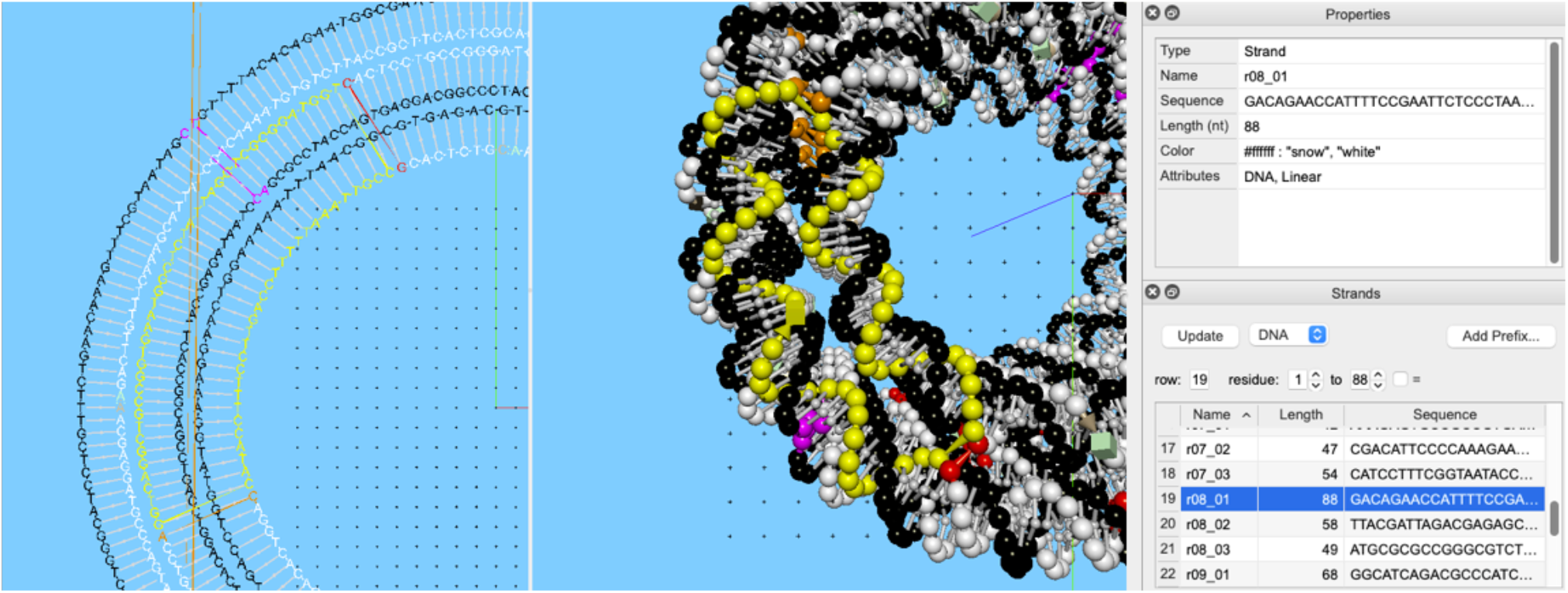
Simultaneous display of Schematic and Design views showing a staple strand that has been selected in the Strand pane (lower right).

In addition to code for performing all major DN design steps, the notebook also offers detailed guidance on navigating inSēquio’s GUI for repositioning the construction plane and performing manual edits. The final steps for constructing the sample design include drawing a single-stranded DNA tether, adding a biotin modification and placing a streptavidin molecule at the barrel’s center (**Figure 10**).

**Figure 10.**
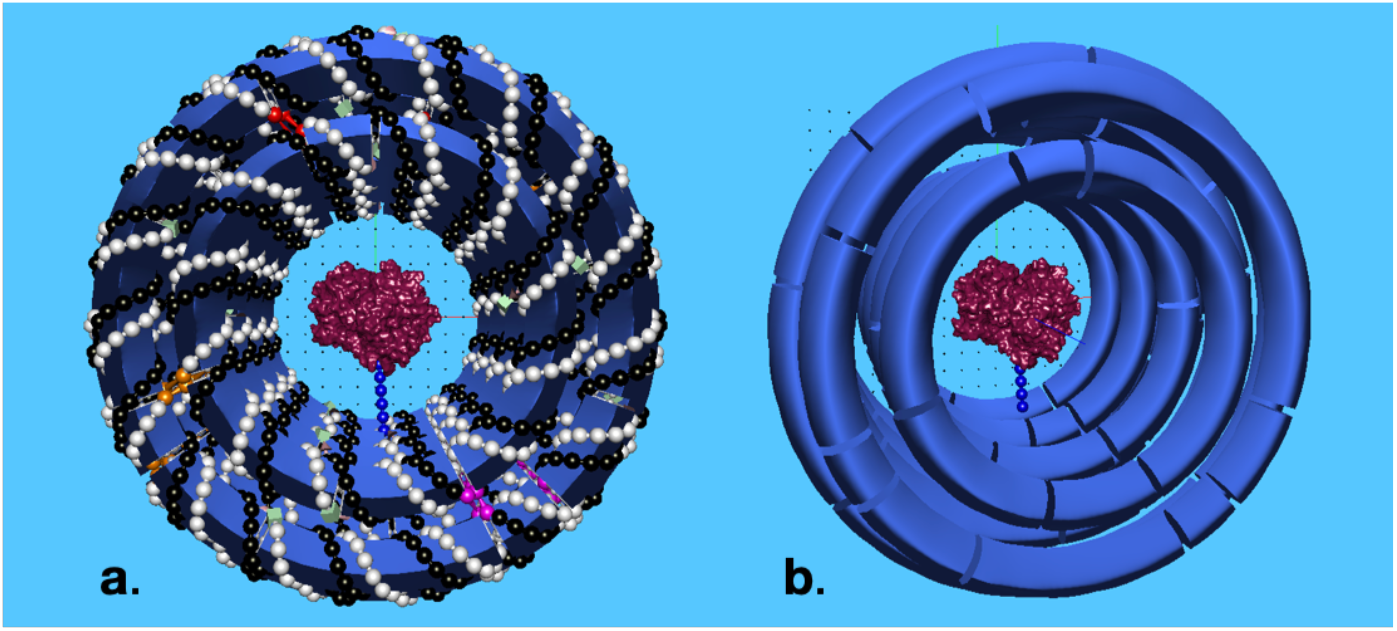
Completed barrel design shown in pipe mode with streptavidin molecule (maroon) atop biotin-tipped single-stranded DNA tether (**a**) with and (**b**) without nucleotides displayed.

It should be noted that even for complex designs, many of the Python methods needed for DN design require only straightforward geometrical or topological calculations, a domain where large-language model (LLM) code generators now excel. For example, the `find_nearest_nucs` function, a simple nearest neighbor search used throughout the notebook, was written entirely by ChatGPT-4 (**Figure S2**). This underscores a significant point: designers with minimal or no Python experience can nevertheless effectively engage in CCD with inSēquio by utilizing sample codes and LLM assistance.

#### A spheroid origami example

Using our CCD approach, we developed a prolate spheroid origami in inSēquio, which is included among the product’s sample designs (**Figure 11a**). It uses an M13mp18 (M13) plasmid as the scaffold and is similar to that of Han et al.^28^ except our design deliberately avoids leaving a scaffold overhang. Since inSēquio draws circular duplexes with an integral number of helical turns, a simple binary search algorithm was used to maximize the number and size of circular duplexes that would approximate a spherical shape and use fewer than M13’s 7249 nucleotides. To prevent strain at the poles, we limited the size of polar rings to a minimum of 128 base pairs, resulting in polar apertures with a radius of ∼11.7 nm. We determined that 23 evenly spaced rings could form a sphere using 7153 scaffold nucleotides, leaving 96 unused. To manage the excess, we incorporated 13 interior-facing scaffold loopouts of 7 nucleotides each, and one of 5 nucleotides (**Figure 11b**). The remaining design steps were executed entirely via Python scripts and the inSēquio API. The crossover and nick selection algorithms used for this design will be integrated into inSēquio in a future release. During the design process, we periodically used inSēquio’s **Export** command to generate oxDNA and all-atom formats for simulation using oxDNA and NAMD, respectively (**Figures 11c and 11d**).

**Figure 11.**
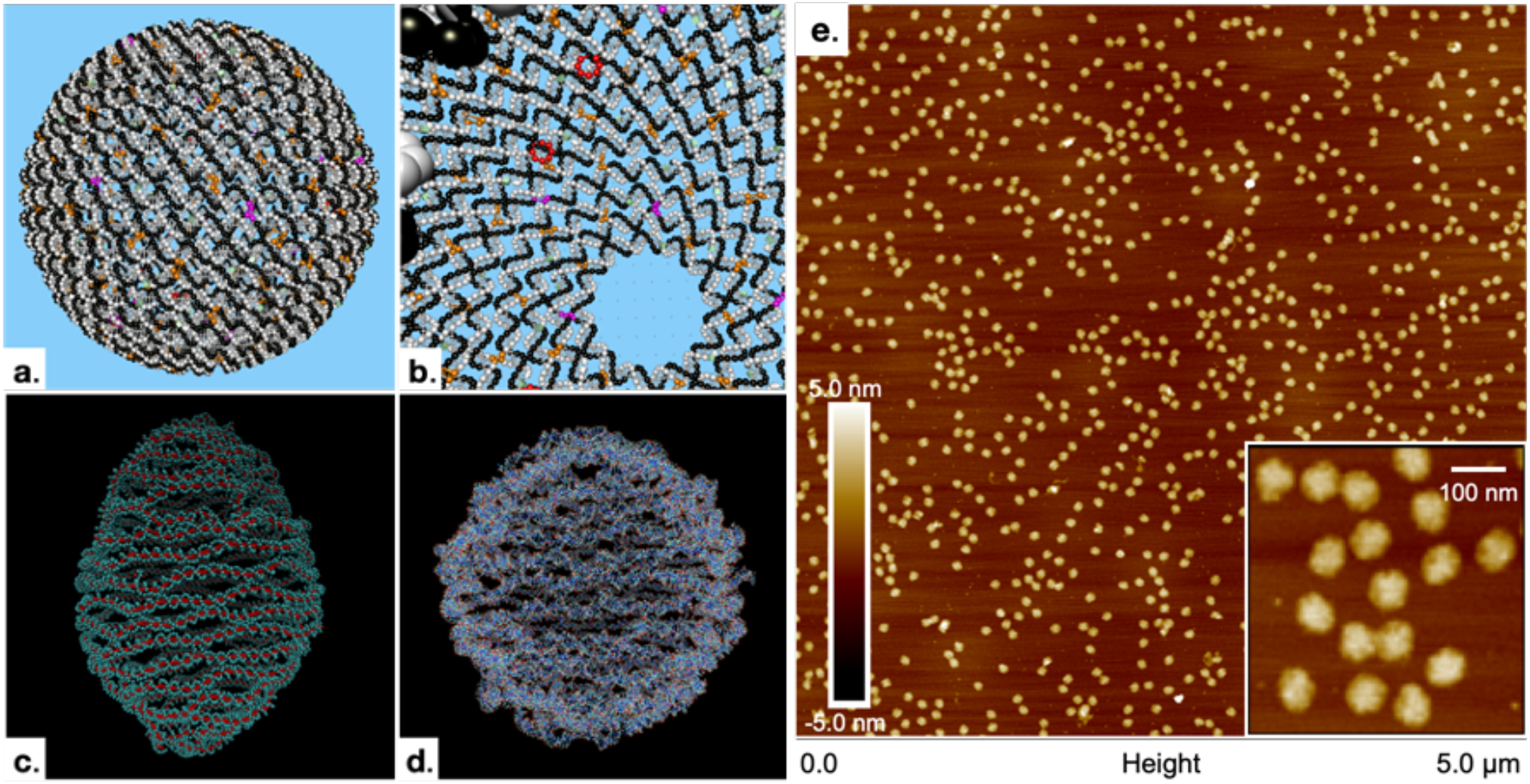
(**a**) A spheroid origami with M13 scaffold created via CCD and shown in inSēquio. (**b**) A view from the interior showing one of the polar apertures and two 7mer loopouts (red). (**c**) Spheroid after conversion to oxDNA format, relaxation, and 50 ns simulation in oxDNA. (**d**) Spheroid after conversion to all-atom PDB format, heating, equilibration, and 5 ns of all-atom, implicit solvent, MD simulation in a CHARMM force field using NAMD. (**e**) An AFM image of the spheroids after synthesis.

### Future directions

#### Cloud-based service offerings

Upcoming releases will leverage the inSēquio Services infrastructure to offer a range of cloud-based services. These services are designed to facilitate compute-intensive tasks and will be available through a usage-based (“metered”) billing model, supported by a commercial cloud provider, such as Amazon Web Services. First will be the integration of all-atom molecular modeling services, which we routinely use in our internal version of the software. Additionally, we have implemented sequence optimization algorithms that minimize undesirable hybridizations (“mismatches”) for both origami (where the scaffold sequence is rotated through all possible start locations) and less constrained designs and these, too, will be offered as metered cloud services.

#### Integrated trajectory viewing

Currently, oxDNA^22^ simulations can be launched and executed via inSēquio Services, however, after users download resulting trajectories, they must use another tool, such as VMD^29^, to view them. In the future, when simulations complete, results will be automatically downloaded after which users will be able to view them directly in inSēquio. Likewise, in-app viewing of all-atom simulation results will also be supported.

#### Lattice view

Several generations of DN researchers have used cadnano^7^ as their primary design software. For square and honeycomb lattice designs, its 2D user interface is elegant and powerful. Because cadnano designs can be imported into inSēquio, we have not prioritized creating such an interface, but we anticipate adding a Lattice view to natively support such designs in the future.

#### Concurrent operation on multiple designs

Currently, the inSēquio UI and scripts are limited to operating on a single design at a time. Ideally, designers using either the UI or the Python API should be able to manipulate multiple designs at once. With this capability, designers using the UI can have multiple designs open at once (e.g., a reference design in a second window) and designers writing scripts can process and analyze multiple design candidates in parallel thereby accelerating exploration of design space. This capability is on our development roadmap, but will likely not appear until the v2.0 release.

#### Configurable duplex dimensions

Our algorithms for drawing duplexes in 3D are internally parameterized by base pair rise, pitch, twist and other common NA geometric variables, however, the interfaces for dynamically manipulating these parameters are limited or nonexistent. Remedying this deficit is a near-term priority.

#### Support for helices

We define a *duplex* as a contiguous run of two or more base pairs between contiguous subsections of two (generally reverse-complementary) nucleotide strands. Duplexes may also be circular, with two circular nucleotide strands of equal length, joined by the same number of contiguous base pairs. We reserve the term *helix* to refer to two or more abutting duplexes, each abutting pair of which is roughly collinear and connected by one strand (not necessarily the same strand for all duplexes in the helix) (see **Figure 12**). This distinction between duplexes and helices is significant in the context of DN design. Duplexes, as more primitive components, have clear definition and extent, and their local geometry is largely independent of surrounding structures, barring any twist or strain from topological neighbors. Helices, in contrast, embody design intent, yet their geometry is governed by surrounding structure. We have developed algorithms to detect helices, and plan to integrate this functionality into inSēquio soon. Currently, however, the application and its API support only creation and identification of duplexes.

**Figure 12.**
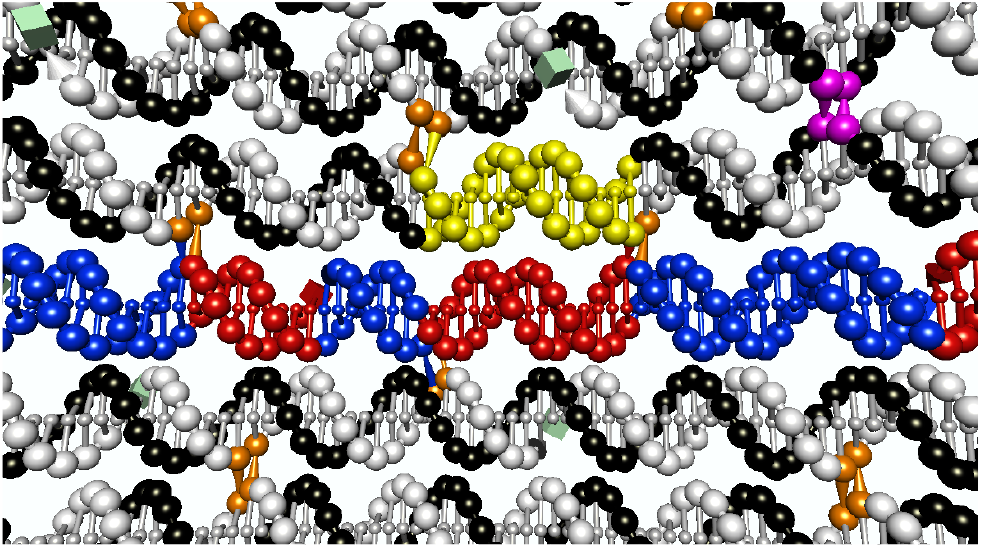
A small patch of nanostructured DNA from a spherical origami design that illustrates the difference between a duplex and helix. A single duplex bound by crossover junctions (orange) is highlighted (yellow). It is part of a circular helix that forms a ring around the sphere. Beneath it is a portion of a similar circular helix with its constituent duplexes alternately colored (blue and red) to emphasize their boundaries formed by crossover junctions or strand nicks.

#### API Enhancements

The Python API currently contains a subset of the functionality provided by the UI. Conversely, there are designs that can only be fully realized through the use of the API (e.g., simultaneous drawing of designs in both the Schematic (2D) and Design (3D) views). More functions will be added to the API to allow programmatic manipulation of designs currently not possible with scripting, including the Find commands, Construction Plane controls, and drawing Molecules. Scene controls such as camera position, view direction and lighting will allow software-driven design animation. Also, the UI will be enhanced to perform as many API-enabled functions as possible. Finally, more complex design logic such as identification of potential crossover sites, helices, and other structurally nontrivial elements, as well as sequence and base pairing optimizations, will be incorporated in inSēquio along with associated API functions to exercise these capabilities where appropriate.

#### Annotations, version control, and provenance

One of the biggest shortcomings of the current version of inSēquio is its lack of support for design annotations. Objects can be named and colored, but otherwise, there is no support for associating notes with a design or its components. The straightforward remedy of providing a notes field for each object is forthcoming, but additional documentation features are desirable, namely, version control and provenance. Users can currently use external version control tools, such as Git or Subversion (SVN), to manage their inSēquio design files, however, a more integrated version control capability may be welcome, especially for users not familiar with software engineering. Our longer-term goal of implementing a provenance system that logs a detailed trace of user interactions will eventually enable several powerful design capabilities. Such traces not only document design history, but they can be converted into scripts (“macros”) that can be saved, optionally augmented with API calls, and executed later to streamline repetitive tasks and improve design efficiency. Particularly useful macros codify desirable features that can be later migrated directly into the product for community-wide efficiency improvements.

## DISCUSSION

The general release of inSēquio, the first commercial fully programmable 3D CAD application specifically created for designing DNs, represents a milestone in the field. It marks the beginning of a transformation in user experience, shifting from the previous necessity of navigating through an assortment of specialized tools to now engaging with a single, comprehensive, and well-supported application – a shift with the potential to significantly enhance productivity for DN designers of all skill levels.

While we have long used prior versions of inSēquio in our research, evolving it into a commercial-grade release demanded substantial effort to achieve the robustness and usability needed for broad distribution. As well, the compilation of a comprehensive User’s Guide, complete with step-by-step tutorials and structured API documentation, and development of the QuickStart wizard each represented significant investments. However, these user support resources, often limited or absent in non-commercial software, significantly enhance the product’s accessibility, which holds the potential to expand the community of DN designers.

Code-centric design (CCD) itself is also becoming more accessible. The advent of large language models (LLMs) capable of generating quality software from prose-based requirements lowers the barrier to employing CCD, especially for designers with little coding experience.

When employed using an interactive 3D CAD environment like inSēquio, CCD offers notable advantages over traditional DN design methods. First, it enhances precision and scalability, which is crucial for managing large, complex DN designs. The reusability of code across different projects provides efficiency gains that quickly outweigh any initial investment in design time. Perhaps most significantly, CCD’s flexibility enables rapid design modifications to meet evolving requirements through straightforward adjustments of parameters and code re-execution.

Finally, the compatibility of CCD with molecular simulation tools like oxDNA and NAMD completes a design cycle that facilitates *in silico* prediction and optimization of DN structures, reducing time-consuming and costly trial-and-error in the laboratory. In conclusion, with its ease of use, support for both freeform and programmatic design, and its ample user support resources, inSēquio is poised to accelerate the use of DNA nanotechnology for a variety of applications, positioning it as an indispensable tool in the field’s future growth and development.

## DATA AVAILABILITY

The M13 sphere design is included with inSēquio. The Jupyter notebook with code for generating the streptavidin barrel is also included.

## ACKNOWLEDGEMENTS

The development of inSēquio would not have been possible without valuable input from the Parabon laboratory team, notably Michael Norton, William Patterson, Michael Gorbet and Hong Zhong. Former Parabon developer Michael Garrahan also made substantial software contributions.

The authors gratefully acknowledge the use of Department of Defense (DoD) high-performance computing resources for the all-atom simulation of the spheroid origami, which were made available under a Small Business Innovation Research contract with the Defense Threat Reduction Agency.

## SUPPLEMENTARY MATERIAL

**Figure S1.**
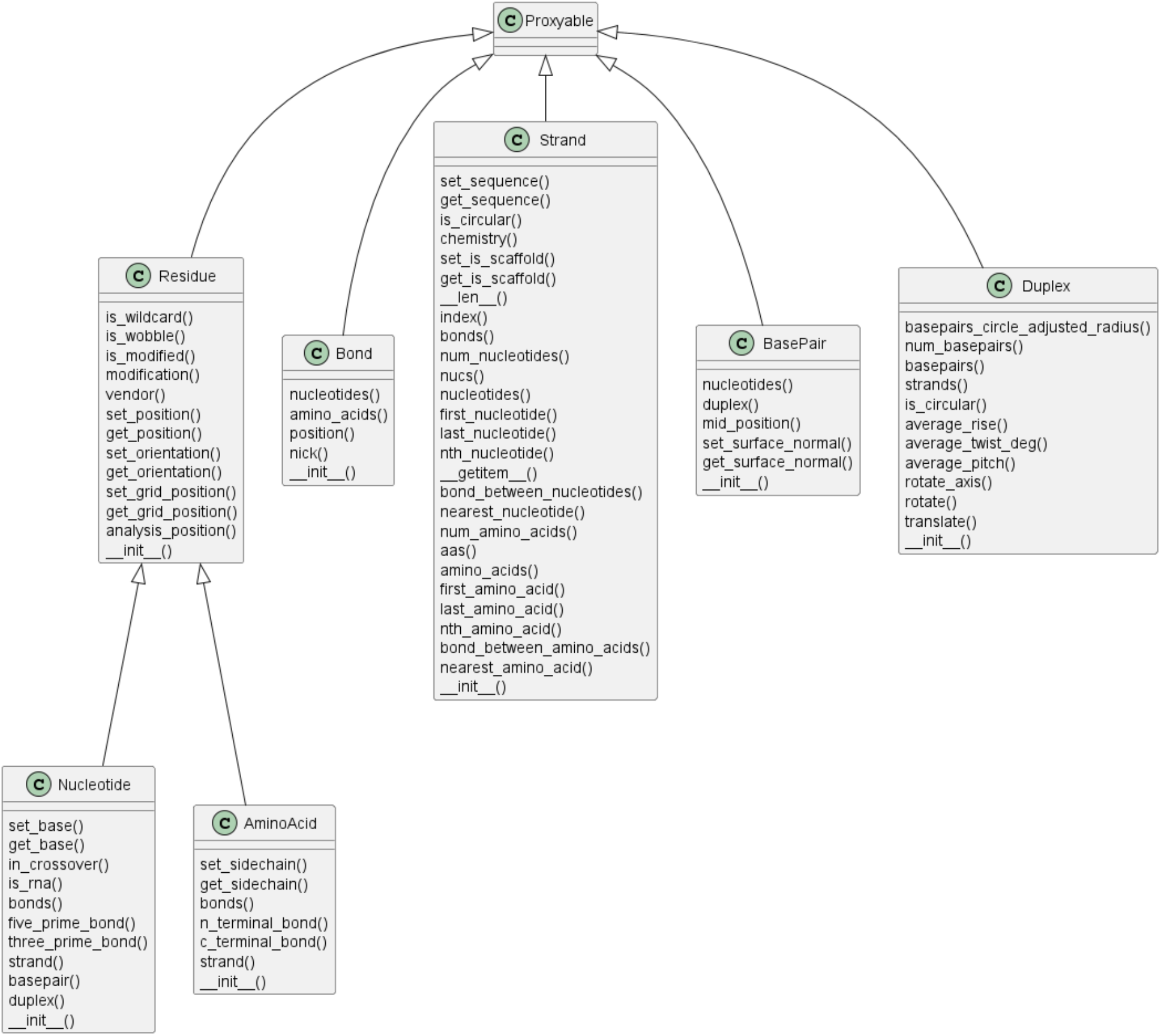
Class hierarchy diagram for the inSēquio API module `insequio.structure`.

**Figure S2.**
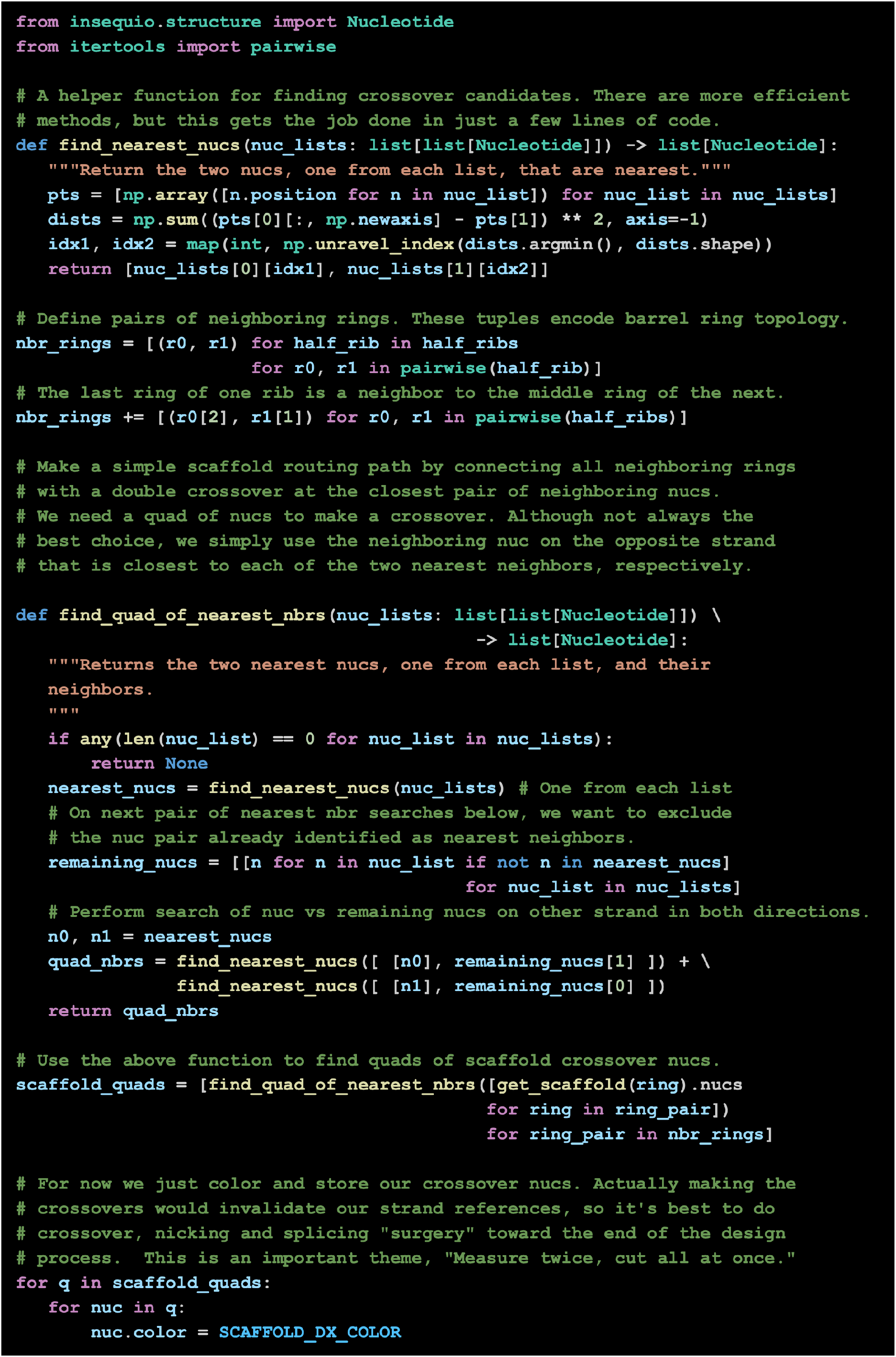
A simple nearest neighbor algorithm for identifying scaffold crossover locations used in the barrel cage sample design.

